# Deterministic Nature of Cellular Position Noise During *C. elegans* Embryogenesis

**DOI:** 10.1101/385609

**Authors:** Xiaoyu Li, Zhiguang Zhao, Weina Xu, Rong Fan, Long Xiao, Xuehua Ma, Zhuo Du

## Abstract

Individuals with identical genotypes exhibit great phenotypic variability known as biological noise, which has broad implications. While molecular-level noise has been extensively studied, in-depth analysis of cellular-level noise is challenging. Here, we present a systems-level quantitative and functional analysis of noise in cellular position during embryogenesis, an important phenotype indicating differentiation and morphogenesis. We show that cellular position noise is deterministic, stringently regulated by intrinsic and extrinsic mechanisms. The noise level is determined by cell lineage identity and is coupled to developmental properties including embryonic localization, cell contact, and left-right symmetry. Cells follow a concordant low-high-low pattern of noise dynamics, and fate specification triggers a global down-regulation of noise that provide a noise-buffering strategy. Noise is stringently regulated throughout embryogenesis, especially during cell division and cell adhesion and gap junctions function to restrict noise. Collectively, our study reveals system properties and regulatory mechanisms of cellular noise control during development.

## INTRODUCTION

Individuals with identical genotypes exhibit great phenotypic variation under homogeneous environmental conditions, a phenomenon known as biological noise (Balazsi et al., 2011). Noise is intrinsic to biological processes that could be both detrimental and beneficial to organismal fitness (Bahar et al., 2006; Burga et al., 2011; Raj et al., 2010). Biological noise is exhibited at various levels from molecular activity to cellular behavior and ultimately organismal function. Noise and its control have been extensively studied at molecular levels, especially the transcriptional (Battich et al., 2015; Elowitz et al., 2002; Faure et al., 2017; Sanchez and Golding, 2013) and post-transcriptional (Fan et al., 2017; Schmiedel et al., 2015) regulation of gene expression noise. However, the properties, regulation, and implications of cellular-level noise during *in vivo* development are poorly understood. Cells are the fundamental structural and functional unit of a complex biological system, connecting molecular activity to organismal function. Therefore, investigating cellular-level noise will shed light on the manifestation and regulation of biological noise at this intermediate and critical level of organization.

*Caenorhabditis elegans* (*C. elegans*) provides an excellent model system to investigate the developmental regulation of cellular noise in metazoans. First, its embryonic and post-embryonic development follows an invariant cell lineage whereby different animals generate the same set of cells in exactly the same way (Sulston and Horvitz, 1977; Sulston et al., 1983). This constancy allows straightforward identification of equivalent cells between animals and the cell-by-cell comparison of cellular behaviors (Murray, 2018). Second, the developmental behaviors of individual *C. elegans* cells are highly stereotypical, including cell division timing and cycle length (Bao et al., 2008; Richards et al., 2013), division axes (Richards et al., 2013), cell fate and differentiation (Du et al., 2014; Murray et al., 2012), cell position (Moore et al., 2013; Richards et al., 2013; Schnabel et al., 2006) and migration patterns (Moore et al., 2013). This predictability makes the analysis of cellular variability (noise) particularly meaningful and biologically relevant. Finally, the transparent cuticle of *C. elegans* combined with advancements in live imaging (Bao et al., 2006; Wu et al., 2013), single-cell tracing (Bao et al., 2006; Jelier et al., 2016; Santella et al., 2014; Santella et al., 2010; Schnabel et al., 1997), and quantitative image analysis (Bischoff and Schnabel, 2006; Moore et al., 2013; Santella et al., 2015; Schnabel et al., 2006; Schnabel et al., 1997) paves the way for systems-level single-cell analysis of cellular noise during *in vivo* development.

For animals (such as *C. elegans*) with a fixed (deterministic) cell lineage and reproducible cellular behavior, a key question is: how is such developmental precision achieved and maintained? Quantifying and analyzing cellular noise would provide a new perspective on this subject. Prior studies have quantified the variability of diverse cellular behaviors during normal *C. elegans* embryogenesis, and reported that despite being stereotypical, cells do exhibit considerable variation (Bao et al., 2008; Richards et al., 2013; Schnabel et al., 1997). However, explicit investigation of the general properties and regulation of noise in cellular phenotypes has not yet been performed. Many key questions about cellular noise remain to be answered, such as: Is cellular noise stochastic or deterministic? Is it coupled to the developmental properties and functions of cells? What regulates cellular noise spatially and temporally?

Here, we performed a systems-level quantitative and functional analysis focused on the variability of 3D cellular position in developing *C. elegans* embryos. Cellular position is a highly regulated and important developmental phenotype that underlies many key processes including cell differentiation, signaling induction, gastrulation, pattern formation, and morphogenesis. A comprehensive investigation of noise in cellular position will shed light on the control of developmental noise at the cellular level. In this study, we reveal general properties, implications, and regulation of cellular position noise during embryogenesis.

## RESULTS

### Quantification of Cellular Position Noise during *C. elegans* Embryogenesis

Cellular position noise (CPN) concerns the variability in a cell’s 3D position among wild-type embryos. To quantify CPN, we determined each cell’s 3D position at minute temporal resolution for each stage of embryogenesis and then measured the variability between embryos grown under identical conditions. To visualize cellular positions, we used nucleus-localized, ubiquitously expressed mCherry to label every nucleus and performed 3D time-lapse imaging (Figure 1A) to record embryogenesis for six hours at 75-second intervals. Based on the fluorescence signal, we identified the 3D position of each cell at each timepoint and traced them to reconstruct the entire cell lineage out to the 350-cell stage using an established method (Bao et al., 2006; Santella et al., 2014; Santella et al., 2010) (Figure 1B). By the 350-cell stage, the embryos have completed all but the last round of cell division and contain over 50% of all cells produced during embryogenesis.

**Figure 1.**
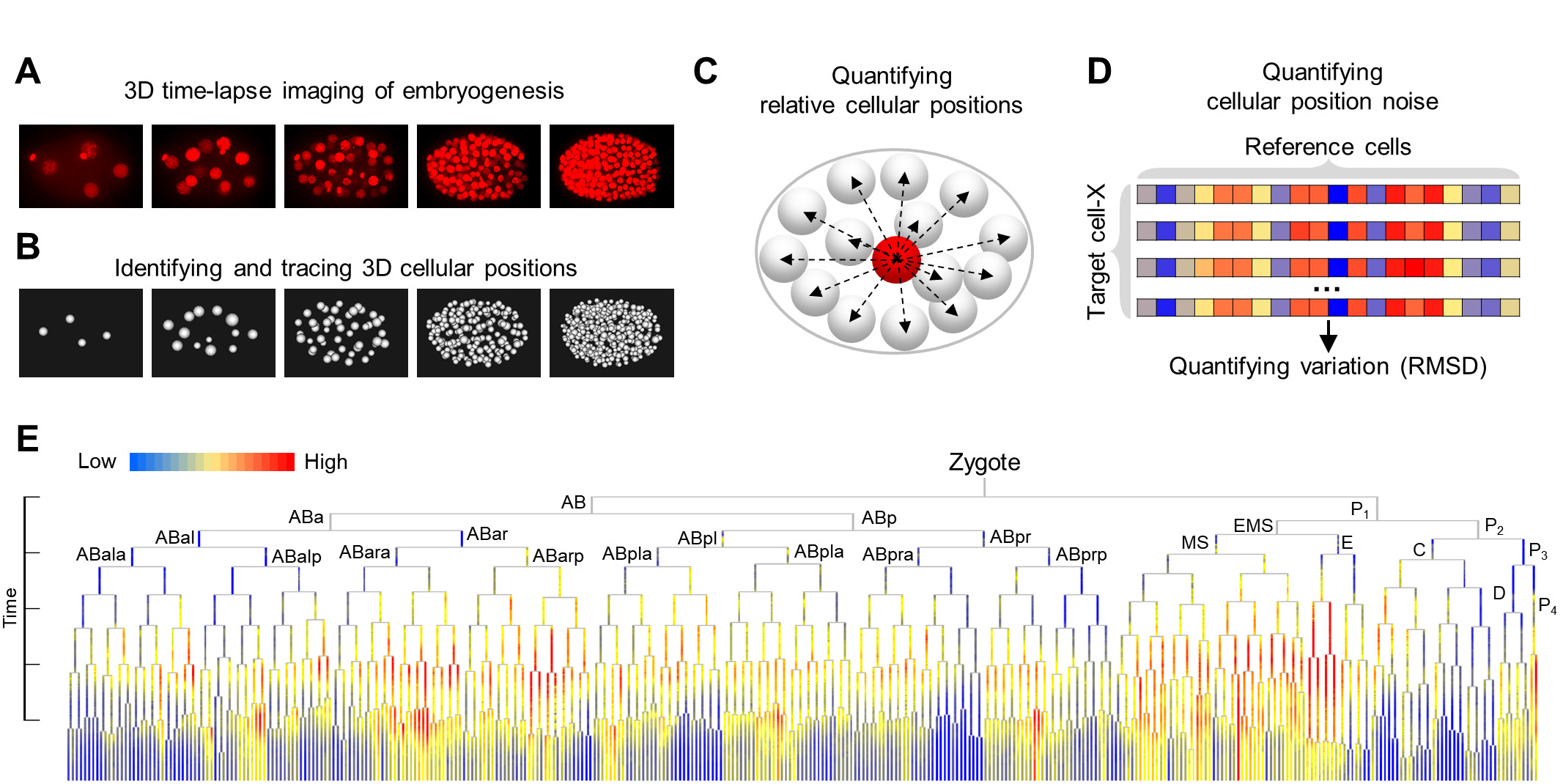
Systematic Quantification of CPN during Embryogenesis. (A) 3D time-lapse imaging of embryogenesis. (B) Cell identification and lineage tracing to determine the position of each cell at each embryogenesis timepoint. See also Table S1. (C) Quantification of relative cellular position as a vector of 3D distances between a target (red) cell and all other cells (white). (D) Determination of CPN by calculating the variation of relative position between wild-type embryos. (E) Tree visualization of CPN levels for every cell during every embryogenesis timepoint. Cells and their lineage relationships are visualized as a tree structure with vertical lines indicating cells and horizontal lines indicating cell divisions. Lengths of vertical lines are proportional to cell cycle length. Cell naming and placement on the tree is according to Sulston nomenclature. See also Figure S1 and Table S1.

For comparing cell positions between embryos, we adopted a relative approach, for representing a given target cell’s position as a vector of geometrical distances from the centroid of its nucleus to those of all other co-existing cells (reference cells) (Figure 1C). We then took CPN as differences in that vector between wild-type embryos (n = 28) for every cell in every aligned timepoint (Figure 1D). Consequently, we systematically quantified CPN at a high temporal resolution across an extended period of embryogenesis, yielding 23,486 data points for 714 cells (Figure 1E and Table S1).

We systematically validated the reproducibility, sensitivity, and robustness of noise measurement (STAR Methods), including the accuracy of cellular position assignment, the robustness and sensitivity of relative quantification of position as a measure of CPN (Figure S1A and Figure S1B), the correctness of time alignment (Figure S1C), the sufficiency of the sample size used (Figure S1 D), the stability of CPN (Figure S1E), and the consistency of CPN measurements across different embryo mounting methods (Figure S1F).

### Intrinsic Regulation: Cell Lineage Identity Determines Cellular Position Noise

Upon obtaining CPN measurements, we first addressed the question: Is CPN stochastic or deterministic? If deterministic, we predicted CPN levels would be significantly associated with the developmental properties of cells. The vast knowledge available concerning *C. elegans* cells in embryogenesis allowed us to correlate CPN with diverse cell properties including cell lineage, cell fate, 3D localization, and left-right symmetry.

Cell lineage is an important cellular property that describes the developmental history of all cells from the zygote. We evaluated whether the lineage history of *C. elegans* cells affects CPN. We classified leaf cells (the traced terminal cells at the 350-cell stage) into 12 groups based on the sub-lineages they derived from, and found that the levels of CPN were sub-lineage-specific (Figure 2A and Table S2). For example, cells from MS and P3 sub-lineages exhibited higher levels of CPN, while cells from the ABpra and C sub-lineages exhibited lower levels. To further elaborate the relationship between CPN and cell lineage, we searched for CPN transition points on the cell lineage tree, defined as ancestral cells that give rise to two sub-lineages with significantly different levels of CPN (Figure 2B). Many transition points were identified (49 points at *p* < 0.05 and 13 points at *p* < 0.001, Wilcoxon Signed-Rank test, Figure 2C and Table S2), suggesting CPN is strongly associated with lineage identity. These transition points partitioned the entire cell lineage line into sub-lineages which each exhibited homogenous levels of CPN (Table S2). Figure 2C shows the 20 sub-lineage groups partitioned by the 13 highly significant transition points. General linear regression analysis revealed that these sub-lineage groups explain a considerable fraction (50.1%) of the variance in CPN levels.

**Figure 2.**
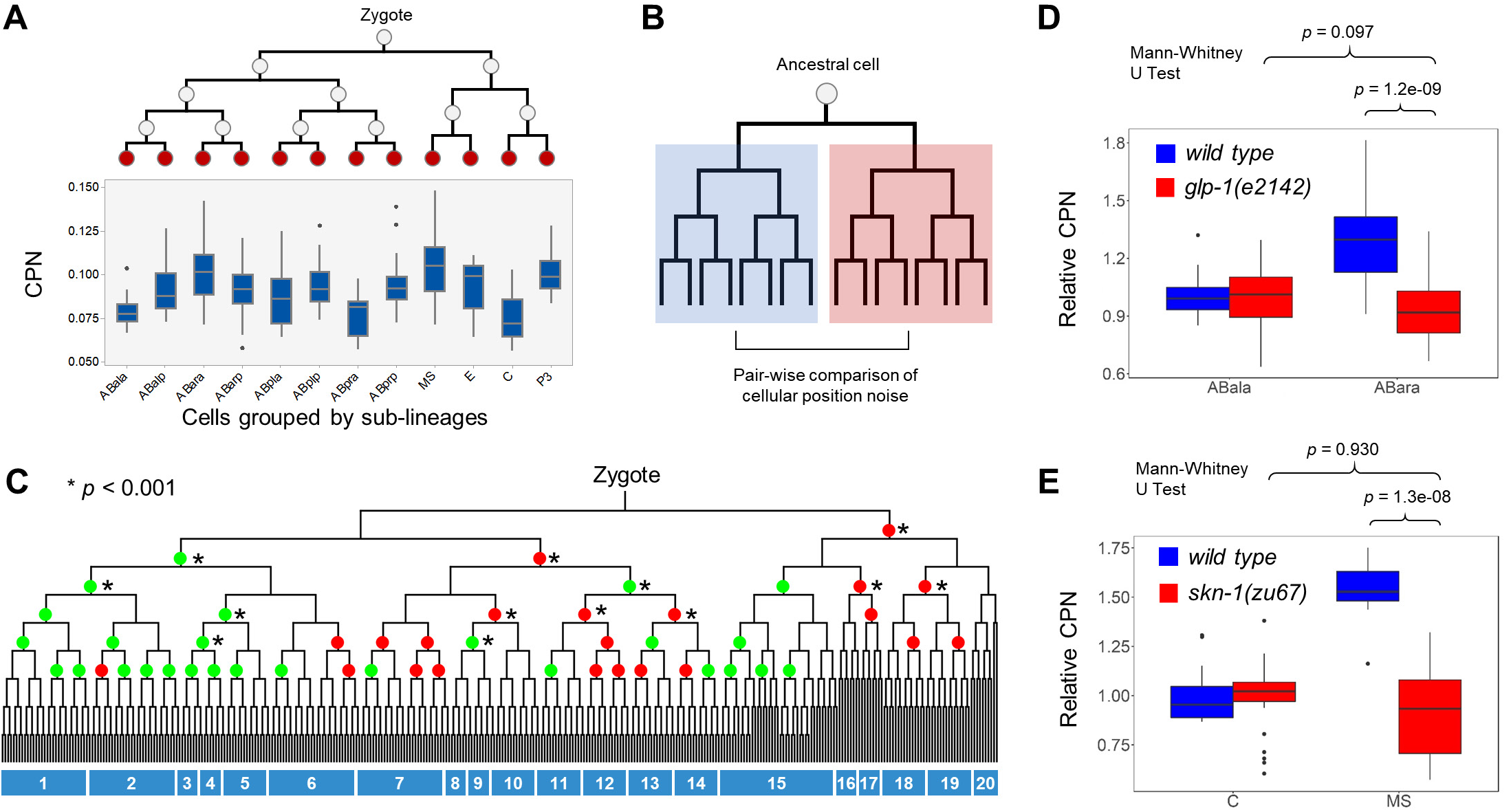
Cell Lineage Identity Determines CPN Levels. (A) Comparison of CPN levels of leaf cells derived from 12 sub-lineages. See also Table S2. (B) The strategy by which CPN transition points were identified. (C) CPN transition points. Circles denote a CPN transition point. Green circles denote those points with a significantly higher CPN of cells from the daughter cell lineage placed on the left side; red circles denote the opposite situation. Stars indicate the transition points with *p* value < 0.001. Numbered rectangles below the tree indicate sub-lineage groups classified by the 13 highly significant transition points. See also Table S2. (D) Changes of CPN levels in *glp-1* mutant embryos. CPN levels of cells from both sub-lineages are normalized to the mean CPN of cells from the ABala sub-lineage. See also Figure S2. (E) Changes of CPN levels in *skn-1* mutant embryos. CPN levels for both sub-lineages are normalized to the mean CPN of cells from the C sublineage. See also Figure S2.

To test whether lineage identity has a causal role in determining CPN levels, we examined how CPN changes when lineage identities were switched. This question was addressed using the ABala and ABara sub-lineages, which exhibited significantly different levels of CPN (Figure 2A). These lineages are distinguished by Notch signaling; loss of function of Notch in the ABara lineage causes its identity to switch to that of ABala (Du et al., 2014; Hutter and Schnabel, 1994; Moskowitz et al., 1994). We expected that if lineage identity underlies CPN differences, CPN levels of the ABala and ABara sublineages in *Notch* mutants would be indistinguishable because the identities of the two sub-lineages are identical. In *glp-1/Notch* embryos (n = 8), we confirmed by tissue marker expression pattern and characteristic programmed cell death (Figure S2A) that the ABara sub-lineage switched to ABala. In *glp-1(e2142)*, we detected a significant reduction of the CPN levels of cells from the ABara sub-lineage as compared to that of *wild type* (Mann-Whitney U Test, *p* = 1.2e-09). In addition, the difference in CPN between the ABala and ABara sub-lineages was no longer significant (Figure 2D, Mann-Whitney U Test, *p* = 0.097). These results are consistent with the expectation that lineage identity determines CPN. We also tested other two sub-lineages (MS and C) and again validated that CPN level changes with lineage identity. These sub-lineages are distinguished by the activity of transcription factor *skn-1/NRF-2*, a master lineage determinant of endo-mesoderm cells whose loss of function induces a lineage identity switch from MS to C (Bowerman et al., 1992; Du et al., 2014). In *skn-1(zu67)*, we found that the CPN of the MS sub-lineage was significantly reduced (n = 11) compared to that of *wild type* (Figure 2E), becoming indistinguishable from that of C (Figure 2E, *p* = 0.930). We also validated that the MS lineage in *skn-1(zu67)* was transformed to that of C based on cell lineage topology (Figure S2B). Together, the above results confirmed a causal role of lineage identity in determining CPN level.

Cell fate is another important developmental property related to the function of cells in the body. In *C. elegans*, cell fate specification is intertwined with but not completely dependent on cell lineage; cells from distinct sub-lineages can differentiate into identical cell types, and those from identical cell lineages can differentiate into distinct cell types. We examined whether different cell types exhibit distinct levels of CPN. Due to technical difficulties, we did not trace the cellular positions through the terminal stage of embryogenesis, but only to the 350-cell stage; nevertheless, by that point, 28% (100) of cells had exited mitosis and differentiated into specific cell types (Table S2). We focused on these cells and compared CPN levels between types; after controlling for lineage identity, we found that that they did not differ significantly (Mood Median Test, *p* = 0.315, Figure 2E and Table S2). Hence, at least by the 350-cell stage, cell fate does not influence CPN.

Taken together, the above correlative and functional data reveal a highly deterministic nature for CPN. These findings suggest that CPN levels are dictated by an intrinsic mechanism that relies on cell lineage but not cell fate.

### Extrinsic Regulation: Cells Similarly Localized and in Physical Contact Exhibit Similar Position Noise

In addition to cell lineage and cell fate, another important developmental property of cells is their 3D localization. Cells occupy specific regions during embryogenesis (Bischoff and Schnabel, 2006; Moore et al., 2013), prompting the question, Does the localization of cells affect CPN? To address this, we clustered cells based on similarity in their relative localization (Figure 3A) and compared CPN levels between clusters. We identified 15 clusters (Figure 3B and Table S3) which exhibited characteristic levels of CPN (Figure 3C). The intra-cluster divergency of CPN was significantly lower than expected (Figure S3A, Mann-Whitney U Test, *p* = 6.55e-100). General linear regression revealed that cell localization explains a considerable fraction (61.7%) of the variance in CPN.

**Figure 3.**
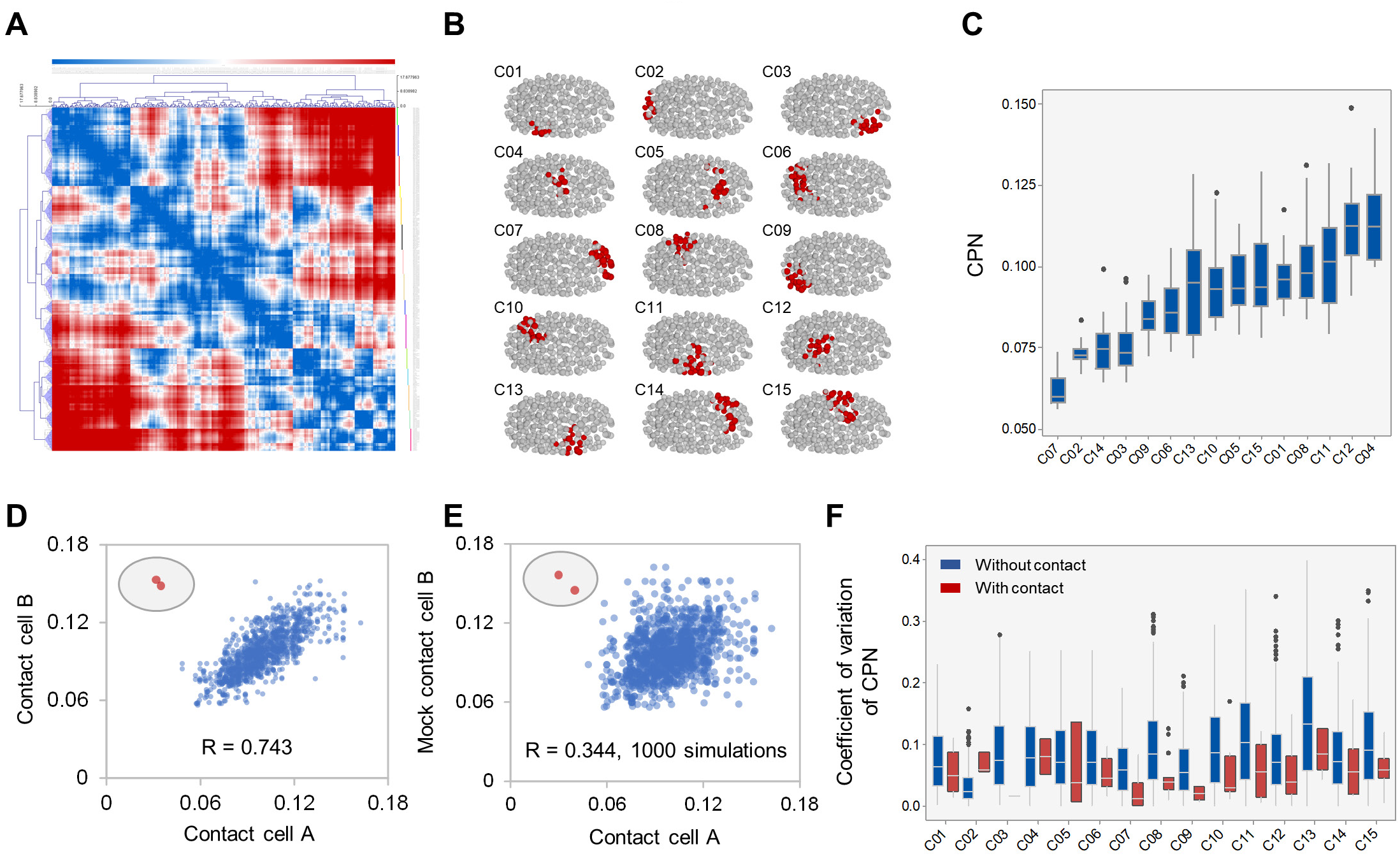
CPN is Associated with Cell Localization and Cell-cell Contact. (A) Clustering of cells with similar localizations. Heatmap shows the hierarchical clustering results of cells based on the pair-wise cell-cell distance. Cell clusters are indicated as colored vertical lines on the right side of the heatmap. See also Table S3. (B) 3D rendering of cells in each cluster. (C) Comparison of CPN levels of cells within each cluster. See also Figure S3. (D-E) Scatter plot showing the correlation of CPN between pairs of contact cells (D) and between randomly selected mock contact cells (E). See also Figure S3 and Table S3. (F) Comparison of divergency of CPN levels for cells in each cell cluster with (red) or without (blue) cell-cell contact. Divergency is measured as the coefficient of variation.

Because both lineage identity and localization were associated with CPN and lineage identity could regulate localization, we examined the influence of each factor while controlling for the influence of the other. We first grouped cells by lineage (Figure 2C) and normalized the CPN level of each cell according to its localization cluster. If cell localization were a confounder of lineage identity, we expected normalization to extinguish the characteristic pattern of CPN in each lineage group. On the contrary, we found that the characteristic levels of CPN were largely preserved (Figure S3B), and CPN values before and after normalization were significantly correlated (Pearson correlation, R = 0.613, *p* = 3.70e-38). These results suggest that CPN is associated with cell lineage even after controlling localization. Similarly, we found continued association of CPN with cell localization after controlling for the influence of lineage identity (Figure S3C, Pearson correlation, R = 0.700, *p* = 1.25e-53). Linear regression revealed that lineage and localization jointly explained 76.8% of CPN variance. Analysis of the relative importance of cell lineage and localization in a linear model showed that in all explained noise variance, the contributions of lineage and localization were generally orthogonal, with localization contributing slightly more than cell lineage (55% vs. 45%).

To further validate the influence of cell localization on CPN, we also applied an alternative strategy for identifying similarly localized cells. In a previous study a Voronoi algorithm was used to infer a reference map describing cell-cell contact during embryogenesis (Chen et al., 2018). By definition, contact cells must be similarly localized. We compared CPN values between contact cells and found that they were significantly correlated (Figure 2D and Table S3, Pearson Correlation, R = 0.743, *p* = 3.76e-195). The correlation coefficient was significantly higher (p = 4.37e-67) than that obtained for mock contact cells (Figure 2E, R = 0.345 ± 0.023, 1,000 simulations). In addition, the divergency of CPN values between contact cells was significantly lower than between mock cells (Figure S3D, Mann-Whitney U Test, *p* = 4.43e-59). This result independently confirmed that cell localization influences CPN.

In addition to having similar localization, contact cells must also physically interact. We therefore examined whether the presence of physical contact further influenced divergency in CPN. We integrated the cell localization clusters described above with the cell contact data and within each localization cluster compared the divergency of CPN between cell pairs with or without cell contact (Table S3). We found that the consistency of CPN levels between contact cells was significantly higher than in those without contact (Figure 3F, Wilcoxon Signed-Rank test, *p* = 3.40e-4), suggesting in addition to similar localization, the presence of physical contact further influences CPN.

Collectively, the above results suggest a potential extrinsic regulation of CPN in which cells with similar 3D localization exhibit similar levels of CPN, especially for those with physical contact.

### Left-Right Symmetric Cells Exhibit Similar Position Noise

As a bilateral animal, the *C. elegans* body plan is generally left-right symmetric. Following embryogenesis and morphogenesis, cells from 30 pairs of sub-lineages (Sulston et al., 1983) are organized in a left-right symmetric pattern (Figure 4A and Table S4). We asked whether these left-right symmetric cell pairs exhibit similar CPN levels. Correlation analysis indicated that the CPN levels of corresponding left and right cells (n = 320) were highly correlated (Figure 4B, Pearson Correlation, R = 0.739, *p* = 1.96e-56). The correlation coefficient was significantly higher (*t*-test, *p* = 4.82e-22) than that obtained for mock left-right symmetric cells (Figure 4C, Pearson Correlation, R = 0.295 ± 0.046, 1,000 simulations). Moreover, the divergency of CPN values between left-right symmetric cells was significantly lower than in mock cells (Figure S4A, Mann-Whitney U Test, *p* = 5.55e-12).

**Figure 4.**
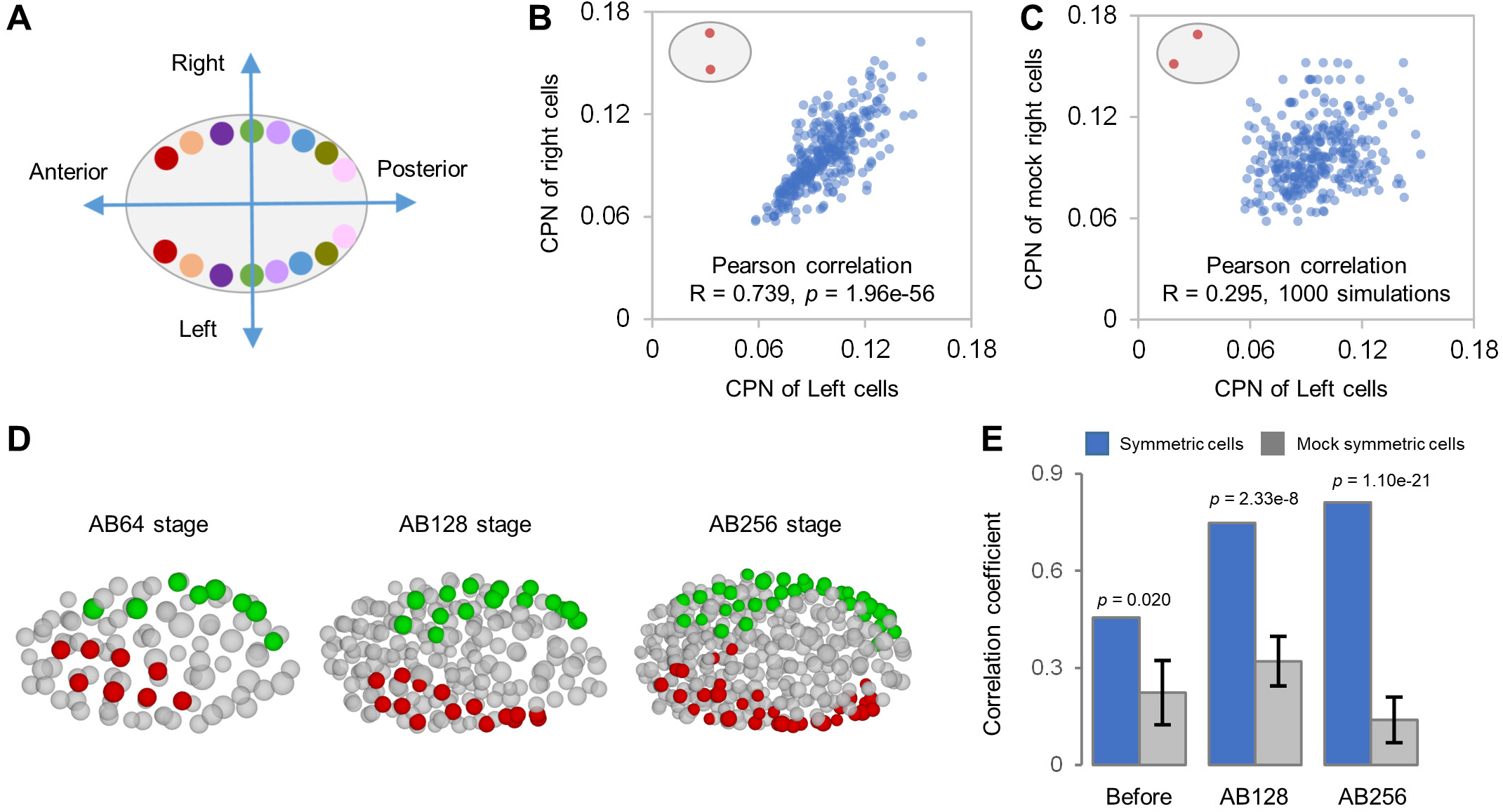
Correlation of CPN Levels between Left-right Symmetric Cells. (A) Schematic representation of left-right symmetric organization of cells. (B-C) Correlation of CPN levels between pairs of left-right symmetric cells (B) and pairs of randomly selected mock symmetric cells (C). See also Figure S4 and Table S4. (D) 3D rendering of pairs of left-right symmetric cells at different developmental stages. Cells from three symmetric lineages (ABplap-ABprap, ABplaaa-ABarpap, and ABplaap-ABpraap) are shown. Red and green represent cells that will localize on the left or right side following embryogenesis, respectively. (E) Comparison of the correlation coefficient of CPN between left-right symmetric cells (blue) and mock symmetric cells (gray) at different developmental stages. *p* value was calculated by *t*-test. See also Figure S4 and Table S4.

Because the establishment of left-right symmetry is progressive (Pohl and Bao, 2010), we examined whether the correlation in CPN levels was specifically coupled to a stage in which symmetry was established. As exemplified in Figure 4D for three pairs of symmetric sub-lineages, the establishment of left-right symmetry is highly time-dependent. We classified the symmetric cell pairs into three categories according to developmental stage (based on the number of AB lineage derived cells, Table S4): before AB128 stage (symmetry not established, 66 left-right cell pairs), AB128 stage (symmetry partially established, 90 cell pairs), and AB256 stage (symmetry well established, 164 cell pairs). If the correlation of CPN between left and right cells were coupled to the establishment of left-right symmetry, we expected the highest correlation and similarity of noise to be present in the AB256 cell stage. Consistently, the highest correlation coefficient (Figure 4E) and lowest divergency (Figure S4B) were obtained for cell pairs at the AB256 cell stage, in which symmetry is well established. In contrast, at developmental stages before AB128, the similarity of CPN between pairs was statistically indistinguishable from that of mock symmetric cells (Figure 4E and Figure S4B). These results therefore suggest that, in addition to cell localization and interactions, the morphogenic organization of cells influences CPN.

### Temporal Regulation: A Concordant Low-High-Low Pattern of Noise Dynamics

Having established the influence of lineage and localization on CPN level, we next sought to answer another question: How do CPN levels change over time? Quantification of global CPN (average CPN across all cells) at each embryogenesis timepoint revealed a smooth and dynamic temporal profile (Table S5). As shown in Figure 5A, this profile follows a low-high-low pattern: that is, during early embryogenesis, global CPN progressively increases with time until mid-embryogenesis (approximately between AB64 to AB128 cell stages); thereafter, the CPN level gradually reduces, and by the 350-cell stage, it reaches a level comparable to the beginning of embryogenesis. This result is consistent with previous visual observations of the variability in cellular position at different developmental stages (Schnabel et al., 1997). To test whether this observed global pattern was an artifact of averaging CPN levels across many cells, we also quantified CPN changes in individual cell developmental tracks. A cell track is defined as following temporally aligned cells within a cell lineage from the ancestral cell to a leaf cell (Figure S5A). We found that individual cells also exhibit a highly concordant pattern of CPN change over time (Figure S5B and Table S5, average Pearson correlation coefficient = 0.719).

**Figure 5.**
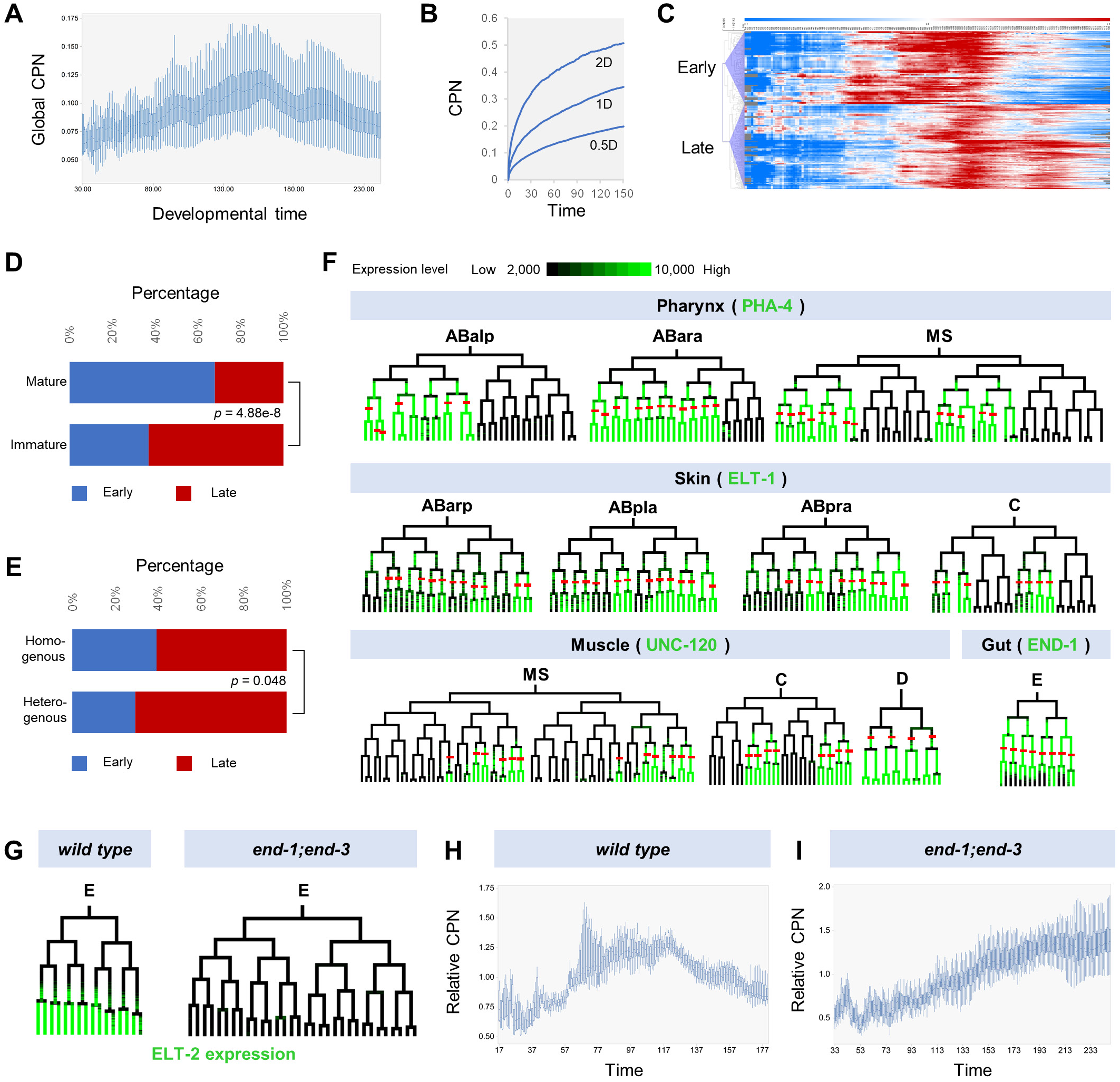
Property and Regulation of Temporal Dynamics of CPN. (A) Changes in global CPN level during every stage of embryogenesis. See also Figure S5 and Table S5. (B) Simulation of CPN level changes over time with various cell displacement distances in one time interval. D is equivalent to the median displacement distance observed in real embryos. (C) Heatmap showing the hierarchical clustering of CPN dynamics. See also Table S5. (D-E) Association of CPN down-regulation with cell differentiation. Stacked bar plot comparing the percentage of cells with earlier (blue) or later (red) CPN down-regulation in cell tracks of mature and immature cells (D) or in cell tracks of homogeneous and heterogeneous progenitors (E). See also Table S5. (F) Expression of fate determinants corresponds with CPN down-regulation. Color coded trees represent single-cell expression of fate determinants in corresponding sub-lineages. Color gradient from black to green depicts expression level from low (<2000 A.U.) to high (>10,000 A.U.). Red lines indicate progenitor cells that give rise to corresponding cell types with down-regulated CPN. See also Figure S5. (G) Abolishment of gut fate specification in *end-1;end-3* embryos. Color coded trees represent the lineal expression pattern of the gut-specific reporter ELT-2 in *wild type* and *end-1;end-3*. (H-I) Changes of CPN dynamics in *end-1;end-3*. Box plot showing the change of global CPN over time for gut cells in *wild type* (H) and *end-1;end-3* (I). See also Figure S5.

We found the pattern of noise change to be tightly aligned to developmental stage rather than the absolute time of development. This was confirmed by comparing CPN dynamic profiles while slowing the global pace of development by reducing the temperature (from 21 °C to 16 °C); this treatment significantly stretched but did not change the pattern of CPN changes (Figure S5C). CPN profiles for the two conditions almost perfectly overlapped when relative time (stage) was used (Figure S5D, Pearson correlation, R = 0.93).

It should be noted that the accumulative nature of cellular position determines that CPN levels over time are not independent values. For example, the CPN of a previous time point will be propagated to the next, and that of a mother cell to its two daughter cells after cell division. Consistently, null models in which the direction of cell displacement was randomized showed a monotone increase of CPN over time (Figure 5B). Therefore, the observed global down-regulation of CPN levels was a consequence of active intervention.

### Fate Specification Triggers Global Down-Regulation of Position Noise

What underlies the global down-regulation of CPN during mid-embryogenesis? A previous study proposed a model in which the blastomere fates could provide spatial codes to its descendant cells, and showed that changes of fate accordingly induce the re-localization of cells (Schnabel et al., 2006). We examined whether cell fate specification and differentiation initiated during mid-embryogenesis underlie the observed down-regulation of CPN.

We first tested whether the timing of CPN down-regulation is associated with the pace of cell differentiation. While the dynamic pattern of CPN was generally concordant across cells (Figure S5B) a closer examination did reveal an asynchrony in the timing of down-regulation. Clustering of temporal changes in CPN by cell track identified two cell groups with differential timing of down-regulation, defined as early and late groups respectively (Figure 5C and Table S5). In the early group, the CPN level peaks and is down-regulated earlier than that of the late group. We then asked: Is this related to cell differentiation? During embryogenesis, the pace of cell differentiation varies across cell lineages; some cells complete mitosis and terminal differentiation earlier than others. We classified leaf cells at the 350-cell stage into two categories, mature and immature, according to whether or not the cell had completed mitosis. We found that a significantly larger portion of cells in the early group were mature than were immature (Figure 5D, Hypergeometric test, *p* = 4.88e-8). We further divided the immature leaf cells into two groups: those that will give rise to identical cell types (homogeneous progenitors) versus those that will give rise to distinct cell types (heterogeneous progenitors). We reasoned that because no further cell differentiation occurs in homogenous progenitors, the differentiated state of these cells would be more mature than those of heterogeneous progenitors. We found that a significantly larger portion of the early cluster were homogeneous progenitors than were heterogeneous progenitors (Figure 5E and Table S5, Hypergeometric test, *p* = 0.048). Together, the above results confirm that CPN down-regulation is associated with cell differentiation.

The overall pace of cell differentiation is determined by the onset of cell differentiation (timing of fate specification) and the pace at which differentiation progresses. We next tested whether the timing of fate specification triggers CPN down-regulation. Specifically, we used the single-cell dynamic protein expression of four well-characterized tissue fate determinants as a readout of the timing of fate specification: PHA-4/FOXA for pharyngeal cells (Horner et al., 1998), ELT-1/GATA for hypodermal cells (Page et al., 1997), UNC-120/SRF for body wall muscle cells (Fukushige et al., 2006), and END-1/GATA for gut cells (Zhu et al., 1997) (Figure 5F). In most cases, the progenitor cells with stable expression of fate determinants showed CPN down-regulation in the same cell or in its daughter cells (Figure 5F and Figure S5E). Only in two out of 101 cases was expression of the fate determinants later than CPN down-regulation. These results suggest that cell fate specification accompanies CPN down-regulation.

To establish a causal role for fate specification in CPN down-regulation, we perturbed fate specification in gut progenitor cells and analyzed the CPN dynamics of their descendants. The specification of gut cells (derived from the E cell lineage) is regulated by a pair of functionally redundant GATA transcription factors, *end-3* and *end-1*; loss of function of both genes abolishes gut fate specification and differentiation (Maduro et al., 2005). We constructed an *end-1;end-3* double mutant and verified that fate specification was abolished in the mutant embryos (Figure 5G), as determined by absence of the characteristic cell lineage pattern and of expression of the gut marker ELT-2. In the mutant, we found that CPN down-regulation was eliminated for cells from the E lineage, and CPN levels accumulated over time (Figure 5H and Figure 5I). Furthermore, the CPN dynamic pattern in other cells was largely unaffected (Figure S5F). This thus validated a causal role for cell fate specification in driving CNP down-regulation.

Together, the above results reveal general properties of the temporal changes of CPN. First, cellular position is highly reproducible in early and late embryos, and CPN dynamics are highly concordant with developmental stage. Second, cell fate specification during mid-embryogenesis triggers global adjustment of cellular position, promoting equivalent cells in different embryos to localize in specific positions, hence reducing noise. This feature provides a noise-buffering strategy that could confer plasticity to embryogenesis (See Discussion).

### Position Noise is Tightly Controlled throughout Embryogenesis, Especially during Cell Division

Because high cellular noise could be detrimental to the robustness of embryogenesis, we next investigated the developmental control of CPN. We first tested whether CPN was subject to tight constraint by comparing observed CPN levels to expected values based on a simulation. We simulated expected CPN by randomizing the directions of cell displacements while keeping the total distance in a cell cycle identical (Figure 6A). The resulting expected CPNs were significantly higher than observed for all cells (Figure 6B, Wilcoxon Signed-Rank test, *p* = 1.49e-118), suggesting CPN is under stringent developmental constraint.

**Figure 6.**
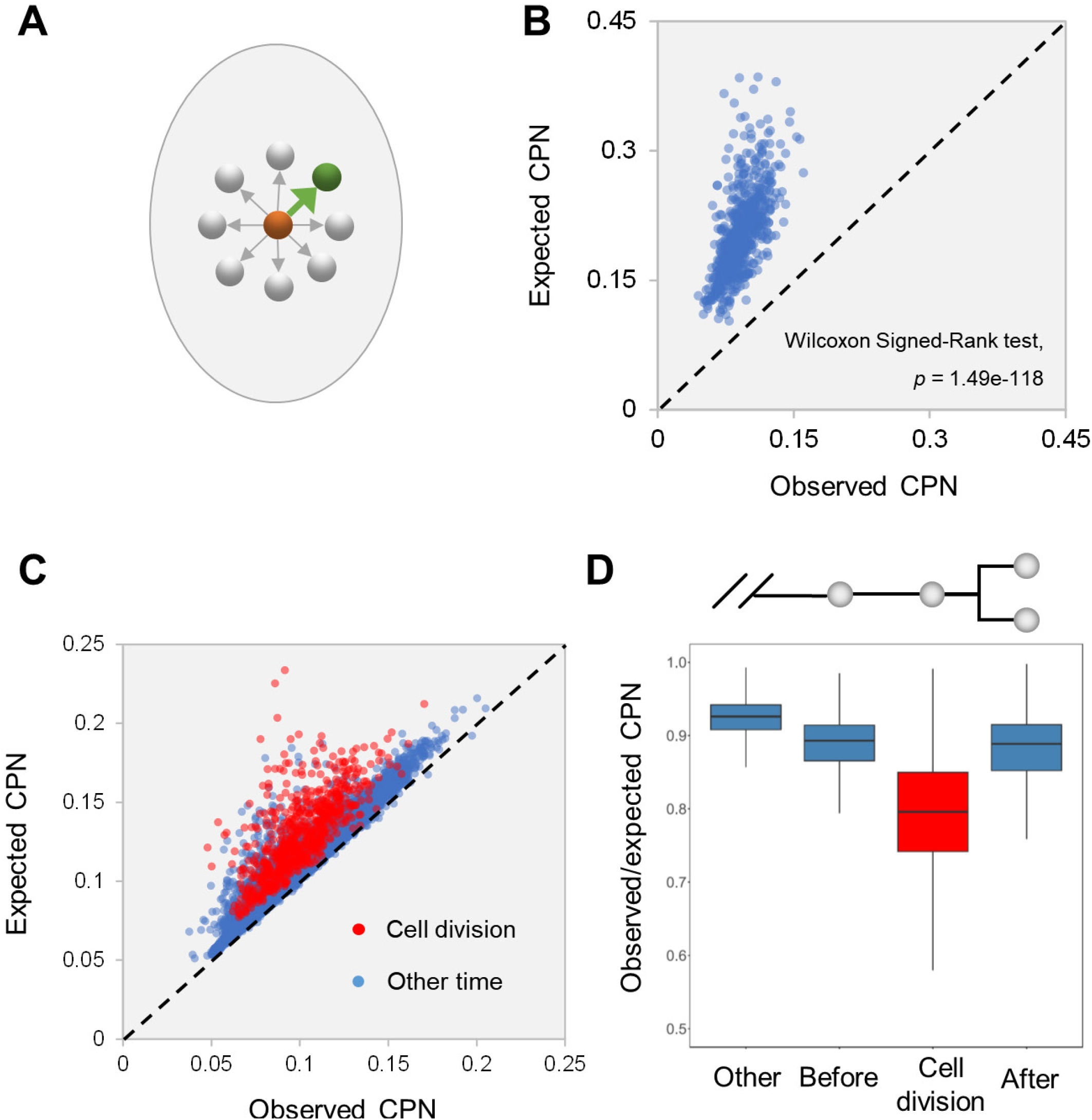
CPN Level is Tightly Controlled during Embryogenesis. (A) Strategy for simulating expected CPN levels. The green arrow indicates the direction and amount of cell displacement observed in a real embryo; gray arrows represent simulated cell displacements. (B-C) Comparison of observed CPN level (X axis) to expected (Y axis) for every cell (B) and for every time interval during embryogenesis (C). Diagonal line indicates equal CPN levels. (D) Comparison of the ratio of observed to expected CPN at different times within a cell cycle.

We next applied the simulation to all cells in two successive timepoints. Once again, the expected CPN was higher than the observed for all cases (Figure 6C), suggesting that cell displacement is directional throughout embryogenesis. Notably, we detected significantly higher constraint on CPN during cell division (Figure 6C and 6D), suggesting that the positioning of daughter cells is tightly regulated during division of most mother cells.

### Cell Adhesion and Gap Junction Function to Restrict Cellular Position Noise

To further elucidate the mechanism underlying restriction of CPN, we analyzed CPN levels in mutants of genes that are potentially implicated in noise control. We reasoned that molecules mediating the cellcell adhesion and gap junctions would be important for restricting random cell displacement and hence reducing noise. To test this possibility, we quantified CPN in mutants of three genes that are important for cell-cell adhesion and gap junctions: *hmp-2/β-catenin* and *jac-1/p120 catenin*, which are components of the cadherin-catenin complex that is required for cell-cell adhesion (Pettitt, 2005), and *inx-2/innexin*, which encodes a gap junction protein known to be expressed during embryogenesis (Altun et al., 2009). In all three mutants, we detected significant increase in CPN levels for the majority of cells (Figure 7A-C and Table S6), suggesting these genes function in noise restriction. Concomitantly, we observed in all mutants that the velocity of cell displacement (average amount cell displacement in a time interval) was also significantly increased as compared to that of *wild type* (Figure 7D-F and Table S6). These results suggest that cell adhesion and gap junctions contribute to noise reduction potentially by restraining the capacity for random cell displacements. However, the characteristic temporal pattern of CPN dynamics was largely preserved in all three mutants (Figure S6A), suggesting these genes do not play essential roles in regulating CPN dynamics.

**Figure 7.**
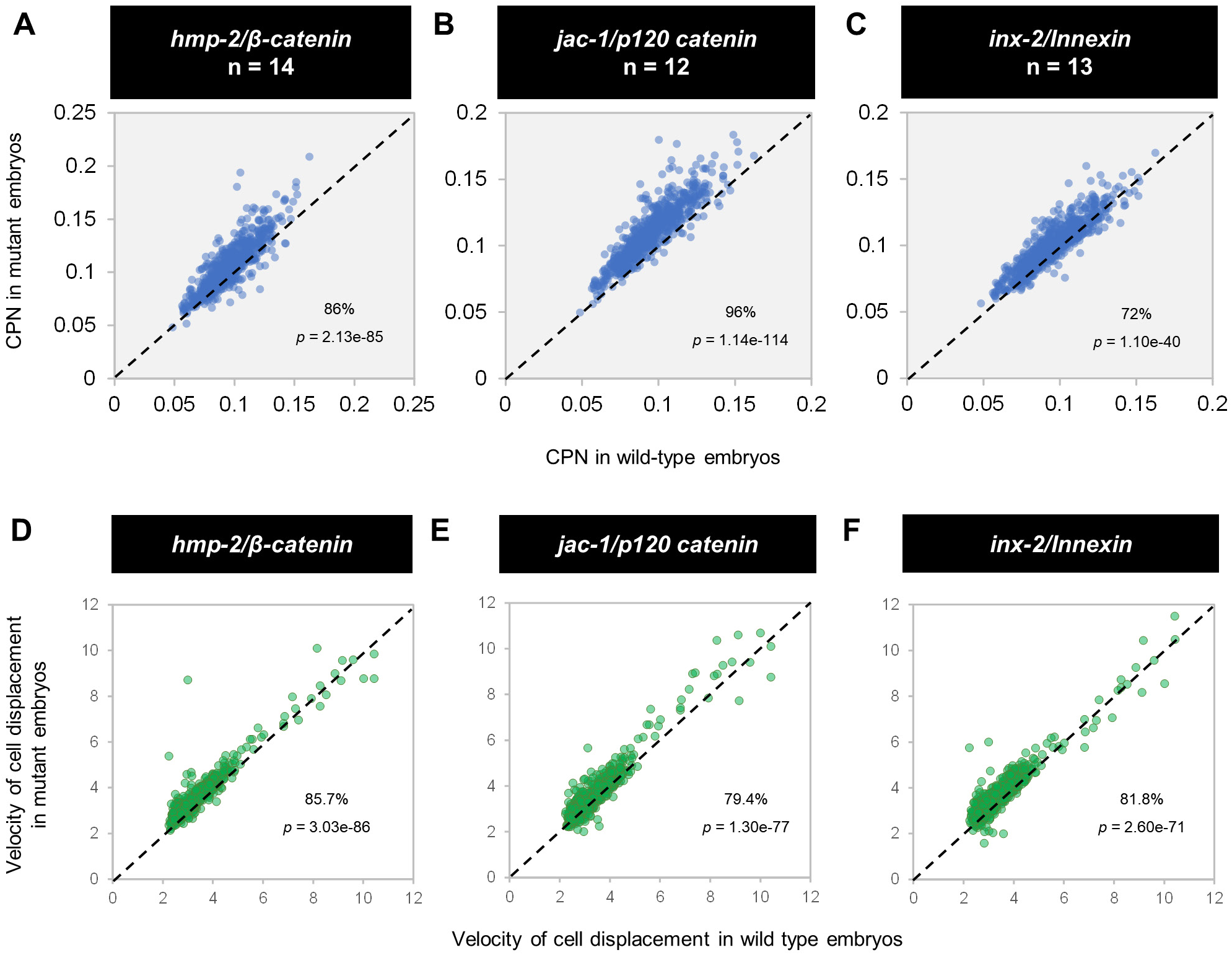
Cell Adhesion and Gap Junctions Restrict CPN Levels. Scatter plots compare CPN levels (A-C) and velocity of cell displacement (D-F) between wild-type and mutant embryos. Velocity is measured as the average distance of cell displacement in one time interval. Diagonal lines represent equal CPN or velocity. See also Table S7.

## DISCUSSION

In this study, we focused on variability in cellular position as an important type of cellular noise and performed a systems-level quantitative and functional analysis. Through live imaging and quantitative single-cell analysis, we have accurately quantified cellular position noise at high temporal resolution throughout an extended portion of embryogenesis (Figure 1). We performed a comprehensive analysis of the properties and developmental control of position noise, with major findings that significantly extend previous knowledge on variability in cellular behaviors (Bao et al., 2008; Richards et al., 2013; Schnabel et al., 1997). We discovered key properties of the level of cellular position noise. Namely, noise level is determined by cell lineage identity (Figure 2) and is coupled to many developmental properties of cells including embryonic localization, cell-cell contact, and left-right symmetry (Figure 3-4). These results reveal a highly deterministic nature of position noise that is potentially regulated by intrinsic and extrinsic mechanisms.

Biological noise has been extensively investigated at the molecular level, especially with regard to gene expression (Battich et al., 2015; Faure et al., 2017). However, due to the technical difficulties involved in quantifying single-cell behaviors and the underlying biological complexities, a systems-level analysis of biological noise at the cellular level has long remained challenging. Focused on cellular position, an important phenotype indicating the differentiation and morphogenic state of cells, our study significantly extends our understanding of the general property of cellular noise during *in vivo* development. In particular, the fact that cellular noise is highly predictable from the developmental properties of cells opens the door to search for intrinsic and extrinsic regulatory factors that determine and controlling the magnitude of cellular noise.

We reveal a highly concordant low-high-low pattern of noise dynamics across diverse cells regardless of their lineage origin, fate, or localization (Figure 5). It seems that embryos can tolerate relatively high systems-wide noise, which will is later relieved by a global down-regulation of noise that is triggered by cell fate specification. We hypothesize that this regulated tuning of noise level could function as a correction mechanism to ensure stereotypical cellular positions in late embryos while also conferring plasticity to embryos. Interestingly, we found in two mutants (*hmp-2* and *inx-2*) that the increased noise relative to *wild type* at earlier stages was partially relieved by this global down-regulation, meaning the comparative noise increase in later embryos is not as severe (Figure S6B). This observation supports that the regulated tuning provides a noise-buffering strategy. A previous study proposed the theory that cell fate provides a positional code that guides cells to localize to specific regions for pattern formation (known as the cell focusing theory) (Schnabel et al., 2006). Here, our data show that a related mechanism could be invoked to reduce and buffer position noise during normal embryogenesis.

Our findings shed light on the systems-wide regulation of position noise. First, we show that position noise is tightly controlled throughout embryogenesis, especially during cell division, to maintain a reproducible cellular configuration in embryos (Figure 6). It is well known that in certain early blastomeres such as EMS (Bei et al., 2002; Sugioka and Bowerman, 2018) and ABar (Walston et al., 2004), the mitotic spindle is tightly regulated during cell division to establish cell asymmetry or to place daughter cells at specific positions for signal receiving. Our data extend previous findings by showing that the division of most cells is stringently regulated to place daughter cells at reproducible positions and that cell movements within a cell cycle are also subject to constraint, although to a lesser extent compared to cell division. Second, our data predict the existence of cell lineage-based intrinsic and cell localization-based extrinsic mechanisms in determining position noise level. As an initial step for characterizing regulators of cellular noise, we identified that proteins mediating cell-cell adhesion and gap junction function to reduce noise by limiting random cell movement (Figure 7). This finding reveals a new developmental function for these molecules (Cox and Hardin, 2004; Simonsen et al., 2014) in maintaining stereotypical cellular position. It will be interesting to test further whether biophysical properties of embryos, distribution of morphogen gradients, or local cell-cell signaling play a role in specifying region-specific noise levels. Together, our study provides insights into the systems properties of and regulatory mechanisms guiding cellular noise during *in vivo* development.

## ACKNOWLEDGEMENTS

This work was supported by grants from the National Natural Science Foundation of China to Z.D. (3177090082 and 31571535), and the “Strategic Priority Research Program” of the Chinese Academy of Sciences to Z.D. (XDB19000000). Some strains were provided by the CGC, which is funded by the National Institutes of Health Office of Research Infrastructure Programs (P40 OD010440).

## AUTHOR CONTRIBUTIONS

X.L. and Z.D. conceived the project and designed the study; W.X., R.F., L.X., and X.M. conducted the experiments and generated the data; X.L., Z.Z., and Z.D. performed the data analysis and statistics; Z.D. wrote the manuscript with input from all authors.

## SUPPLEMENTAL INFORMATION

Supplemental Information includes six figures and six tables.

### Supplemental Figure Legends

**Figure S1.**
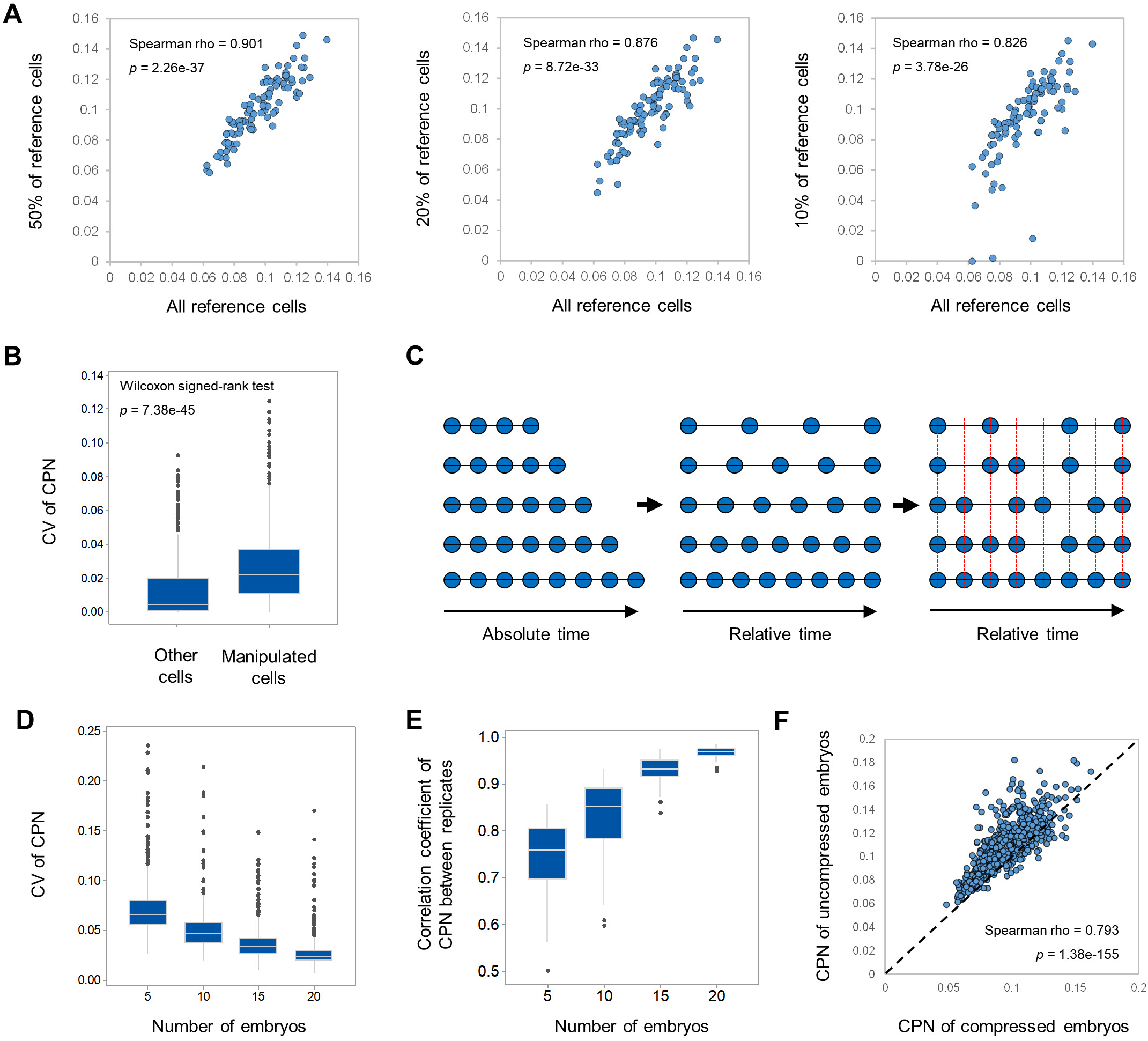
Quality assessment of CPN measurement. (A) Influence of the composition of reference cells on CPN levels. Scatter plots show the correlation of CPN level calculated using all reference cells to that using a subset of reference cells. (B) Assessment of the sensitivity of CPN measurement. Box plot compares the influence of cellular position changes on the CPN levels of position-switched and unaffected cells. (C) Strategy for time alignment. Details are described in the STAR Methods. (D) Influence of embryo number on CPN levels. Box plot shows the coefficient of variation of CPN levels for calculations using a small number of embryos and that using all embryos. (E) Stability of CPN levels. Box plot shows the correlation coefficient between CPN levels in replicate samples using different numbers of embryos. (F) Influence of embryo mounting methods on CPN levels. Scatter plot shows correlation of CPN measurements between compressed (X-axis) and uncompressed embryos (Y-axis).

**Figure S2.**
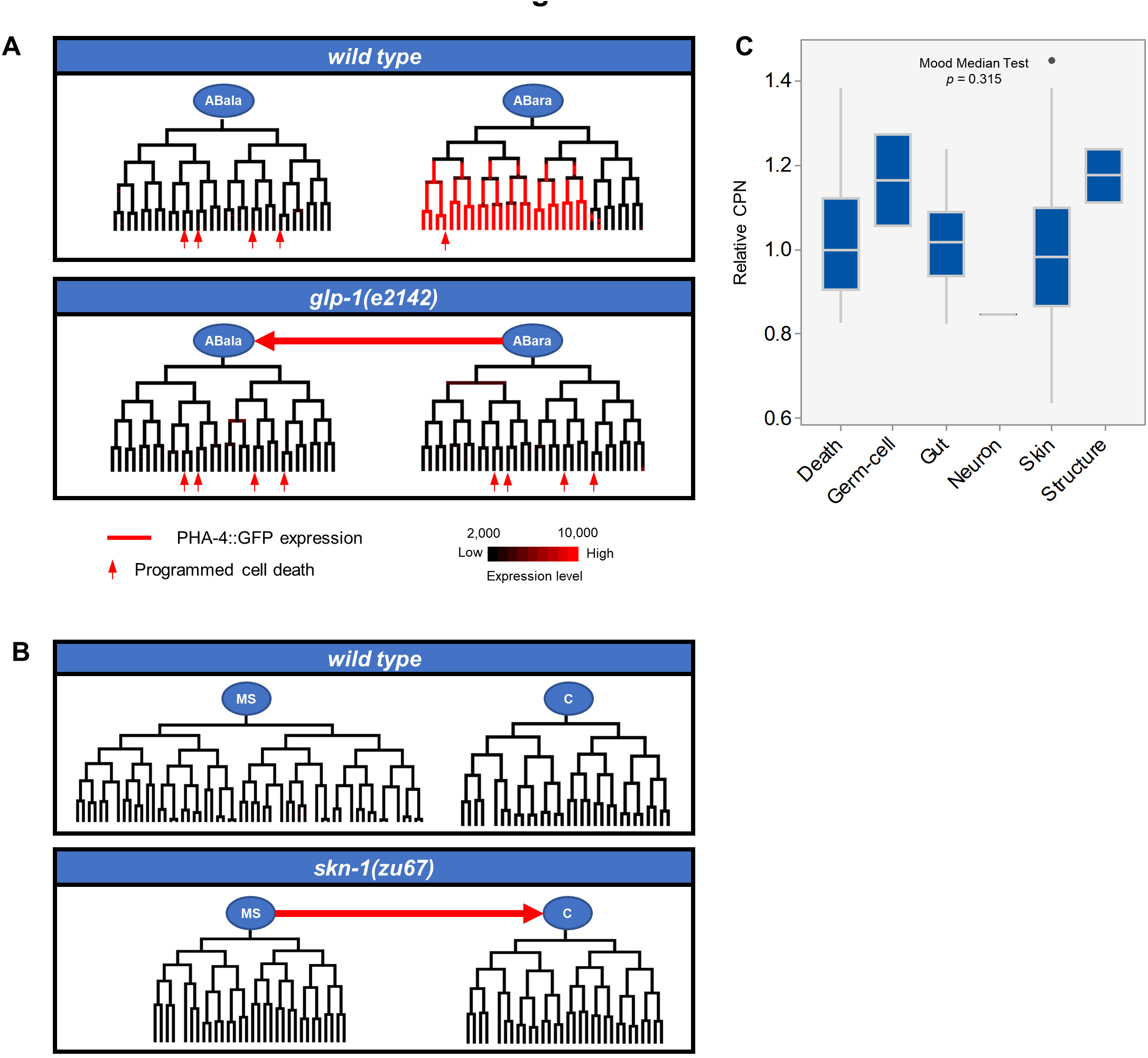
Cell lineage identity but not cell fate determines CPN level. (A) Switch of ABara sub-lineage identity to that of ABara in *glp-1*. The color coded tree shows the traced cell lineage and cellular expression pattern of an ABara-sub-lineage marker (PHA-4∷GFP). Arrows indicate programmed cell death. (B) Switch of MS sub-lineage identity to that of C in *skn-1*. (C) Comparison of CPN levels of cell types. CPN levels of cells are normalized by the average CPN for their sub-lineages.

**Figure S3.**
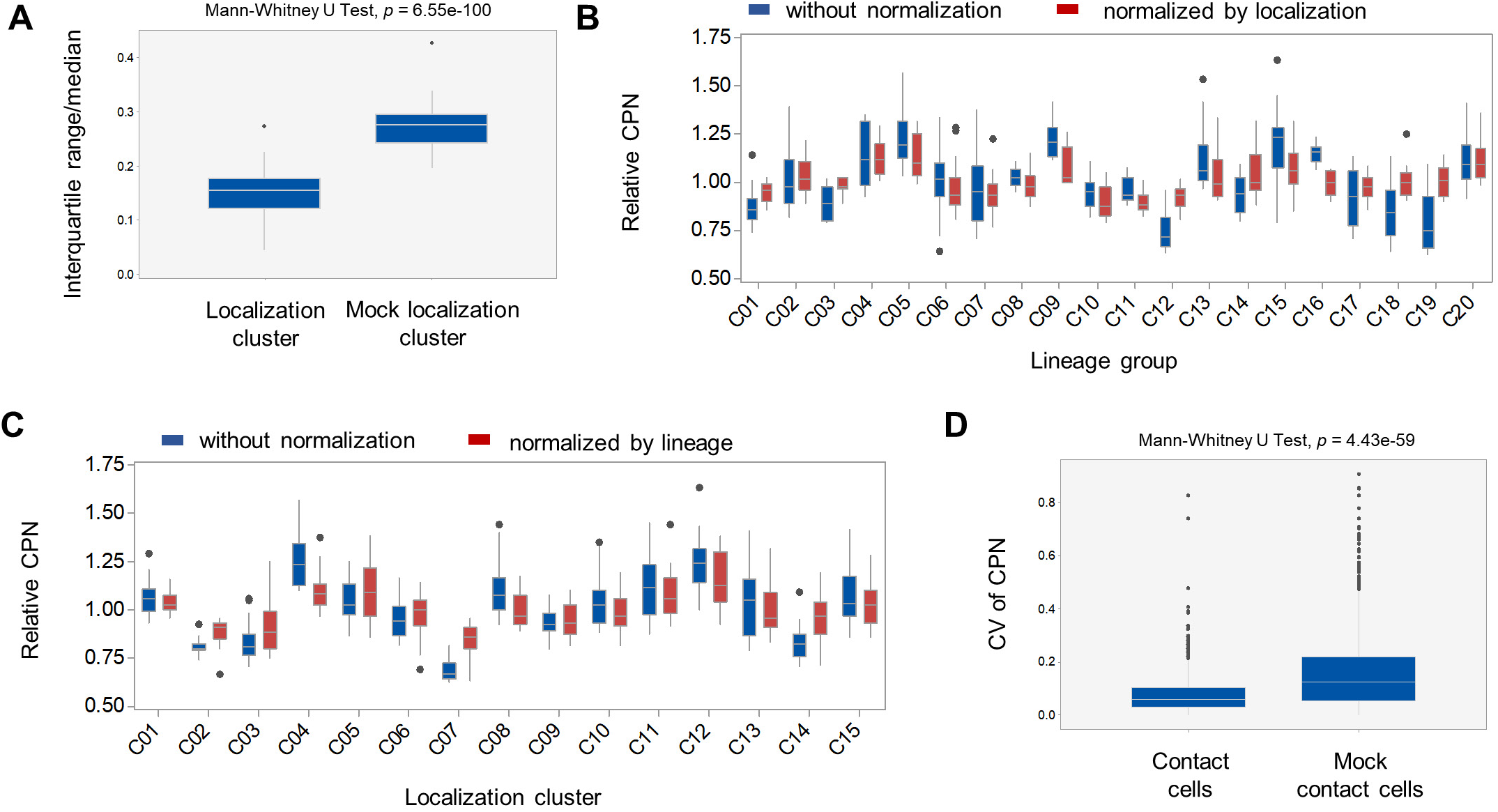
CPN level is associated with cell localization and cell contact. (A) Cells with similar localization exhibit smaller divergency in CPN level. Box plot shows the divergency of CPN levels (measured as interquartile range/median) in real and mock localization clusters. (B) Influence of cell lineage identity on CPN after considering localization. Box plot shows unnormalized (blue) and normalized (red) relative CPN levels for cells from different lineage groups (Figure 2C). Cell CPN was normalized via dividing by the average CPN of its cell lineage group. (C) Influence of cell localization on CPN after considering cell lineage identity. Box plot shows unnormalized (blue) and normalized (red) relative CPN levels of cells from different localization clusters (Figure 3B). Relative CPN was calculated by dividing the CPN of each cell by the mean CPN level. Cell CPN was normalized via dividing by the average CPN of its localization group. (D) Contact cells exhibit smaller divergency in CPN levels. Box plot shows the coefficient of variation of CPN levels between cell pairs from real and mock contact cells.

**Figure S4.**
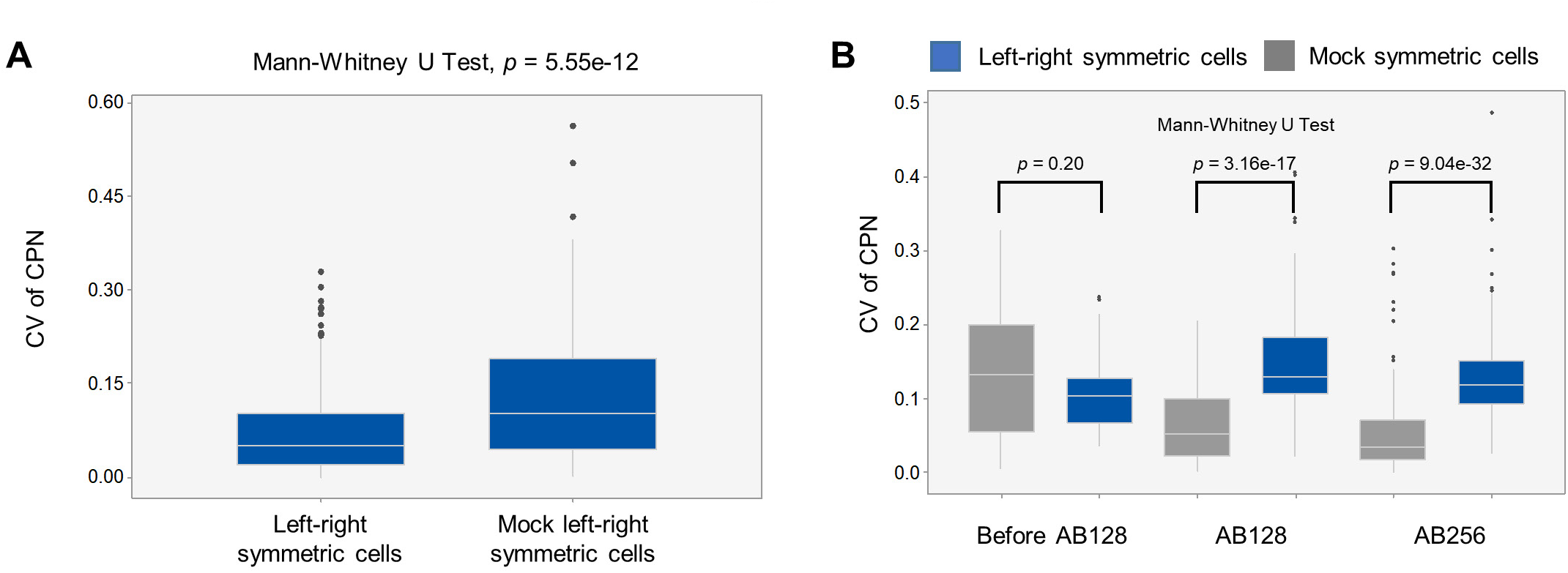
Correlation of CPN levels between left-right symmetric cells. (A) Left-right symmetric cells exhibit smaller variation in CPN levels. Box plot shows the coefficient of variation of CPN levels in real and mock left-right symmetric pairs. (B) Correlation of CPN between left-right symmetric cells is associated with the establishment of symmetry. Box plot compares the deviation of CPN levels between real and mock symmetric cells at different developmental stages.

**Figure S5.**
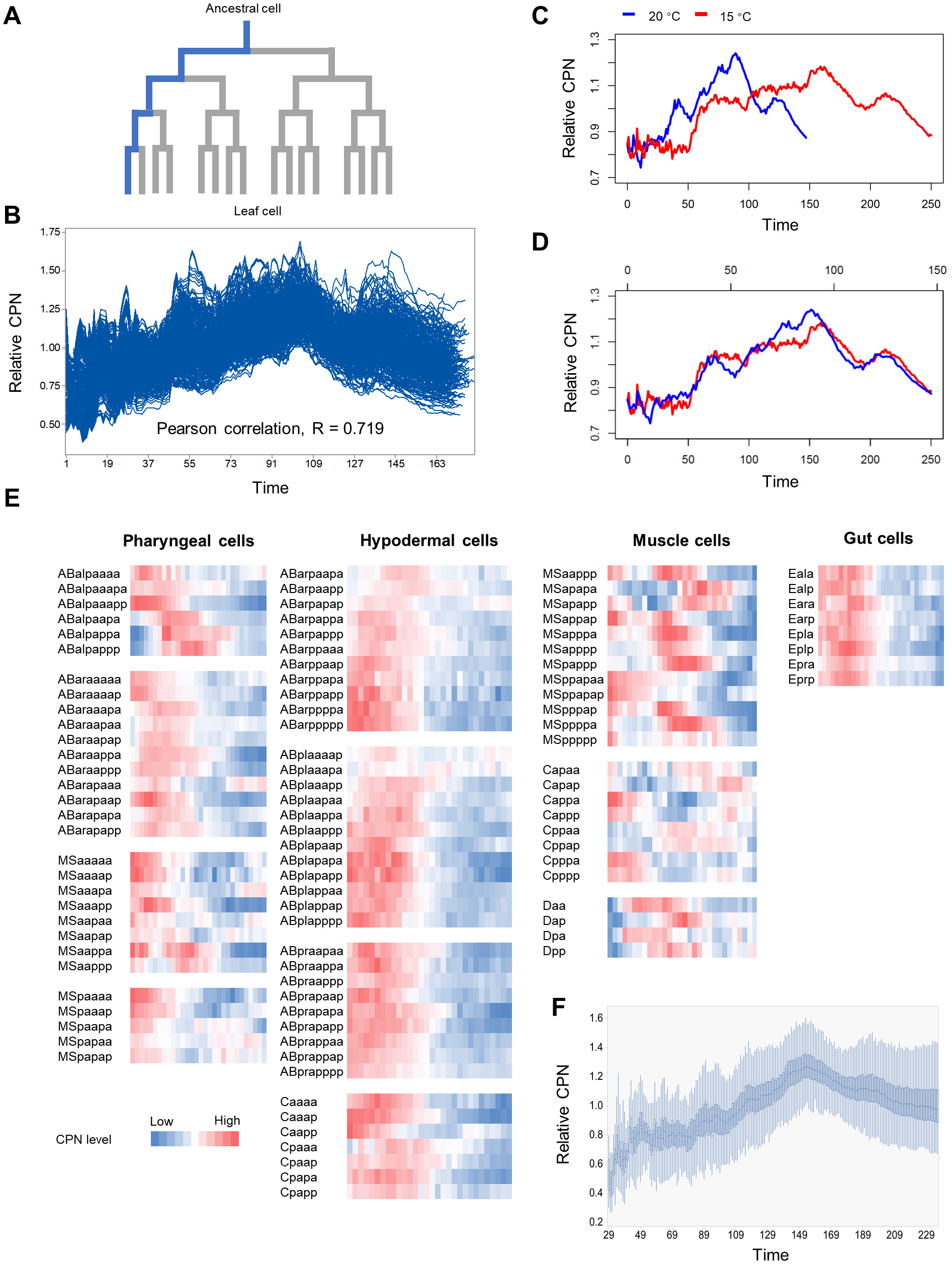
Temporal dynamics of CPN. (A) Definition of cell tracks for each leaf cells. (B) Dynamics of relative CPN levels in each cell track during embryogenesis. Relative CPN was calculated by dividing CPN level at each time point by the mean CPN of a cell track. (C) Comparison of CPN dynamics for embryos cultured at different temperatures. (D) Alignment of CPN dynamics by relative time. (E). Changes in CPN levels during the cell cycle of individual cells with different cell fates. Color gradient from blue to red represents CPN levels from low to high. (F) Dynamics of global CPN levels of non-gut cells in *end-1;end-3* embryos.

**Figure S6.**
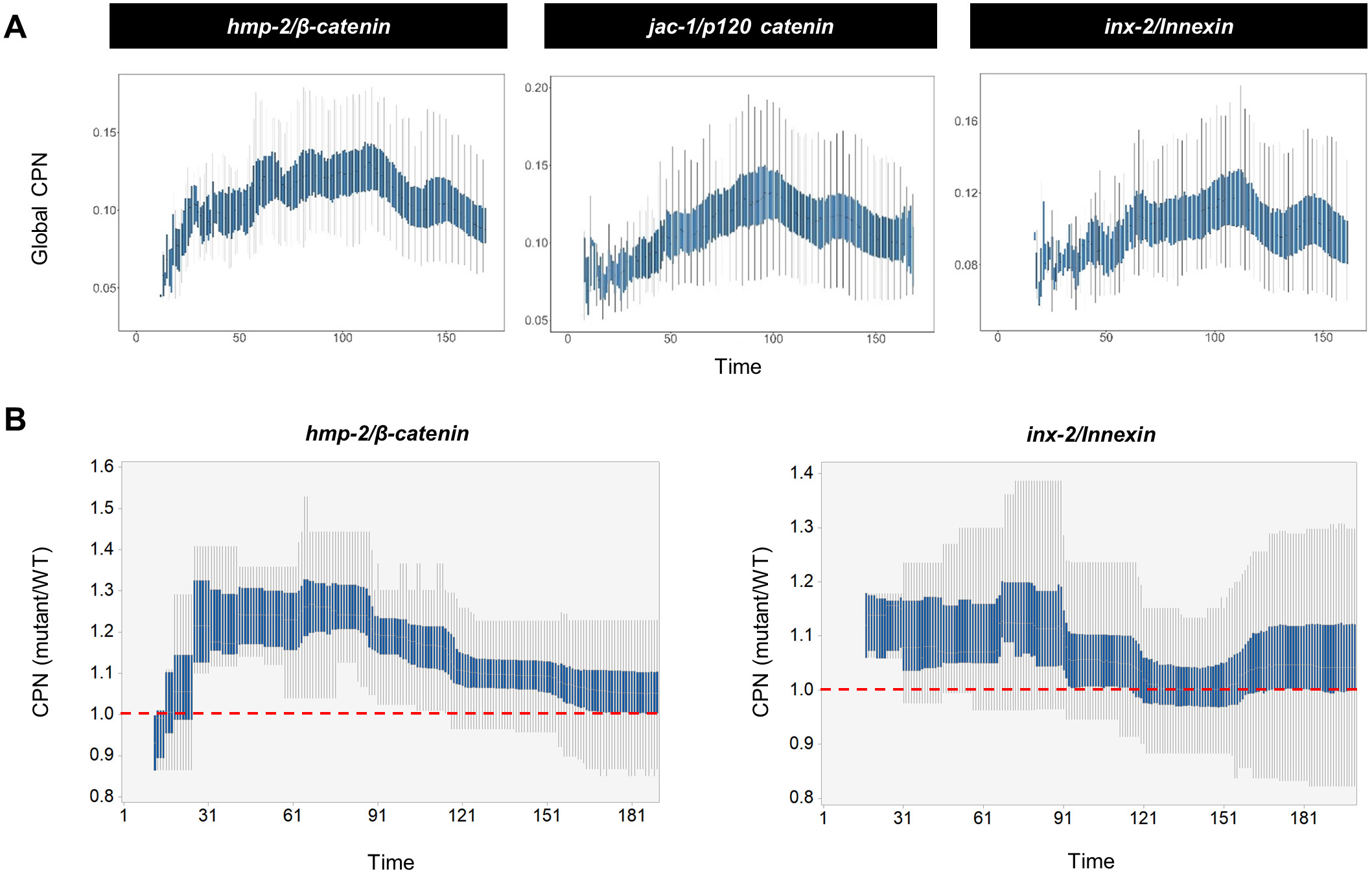
Temporal Dynamics of CPN in Cell Adhesion and Gap Junction Mutant Embryos. (A) Dynamics of global CPN in *hmp-2, jac-1*, and *inx-2* embryos. Box plot shows average CPN across all cells at each developmental time during embryogenesis. (B) Change of CPN over time in *hmp-2* and *inx-2* embryos relative to *wild type*. For ease of comparison, a cell’s average CPN level was used to calculate its relative CPN (measured as CPN of mutant/CPN of WT for equivalent cells) at different times in embryogenesis.

**Table S1. Quantification of cellular position and noise**

1^st^ Sheet lists the position of each cell at each time point in 28 wild-type embryos.

2^nd^ Sheet lists CPN levels of each cell at each time point.

**Table S2. CPN levels of cells from different sub-lineages**

1^st^ Sheet lists CPN levels of cells from different sub-lineages.

2^nd^ Sheet lists the identified CPN transition points.

3^rd^ Sheet lists CPN levels of cells from cell lineage groups classified by transition point with *p* < 0.001.

4^th^ Sheet lists CPN levels of different cell types.

**Table S3. CPN levels of cells with similar localization**

1^st^ Sheet lists CPN levels of cells in each localization cluster.

2^nd^ Sheet lists CPN levels of contact cells.

3^rd^ Sheet lists CPN levels of cells in each localization cluster with or without cell contact.

**Table S4. CPN levels of left-right symmetric cells**

1^st^ Sheet lists CPN levels of left-right symmetric cells at different developmental stages.

**Table S5. CPN dynamics**

1^st^ Sheet lists CPN levels of cells at each developmental time.

2^nd^ Sheet lists CPN levels of each cell track.

3^rd^ Sheet lists the classification of cell tracks according to noise dynamics and differentiation state.

**Table S6. Comparison of CPN levels and cell displacement velocity in wild-type and mutant embryos**

1^st^ Sheet lists CPN levels of cells in wild-type and mutant embryos.

2^nd^ Sheet lists velocity of cell displacement in wild type and mutant embryos.

## METHODS

### *C. elegans* Strains and Culture

Some *C. elegans* strains used in this study came from the Caenorhabditis Genetics Center. All strains were cultured under standard laboratory conditions on nematode growth media plates seeded with OP50 bacteria in incubators at a constant temperature. Most strains were maintained at 21 °C; temperature-sensitive mutants were maintained at 15 °C and switched to 25 °C for at least eight hours before the experiment.

### CRISPR/Cas9-based Gene Knockout and Knockin

Knockout of *end-3* gene: The *end-1* and *end-3* genes are tightly linked on chromosome V. To bypass the inherent difficulty of crossing, we constructed the *end-1;end-3* double mutant using CRISPR/Cas9-based gene editing to knockout the *end-3* gene in *end-1* mutants. Two single-guide RNA (sgRNA) sequences targeting the N- and C-terminals of *end-3* were designed using the CRISPR Design Tool (http://crispr.mit.edu/) and cloned into a pDD162 plasmid by circular PCR using a previously-published protocol (Paix et al., 2014) to construct the Cas9-sgRNA plasmid (end-3sg). This plasmid and a reporter plasmid pRF4 (inducing a dominant roller phenotype) were purified with the PureLink PCR Micro kit (Invitrogen) and injected into the gonads of worms at a final concentration of 50 ng/μL for each plasmid. Single-worm PCR was performed on individual F1 animals with a roller phenotype to detect gene knockout events. Progeny of worms with a large gene deletion were selected to obtain homozygous knockout animals. We obtained a knockout strain with a 931 bp deletion (chr V: 14,014,806-14,015,796, WS235/cel11) and a 9 bp insertion (TTTCAATGA) in the *end-3* gene.

Knockin of mNeonGreen in the *end-1* gene: To visualize endogenous END-1 protein expression, we fused a codon-optimized coding sequence of *mNeonGreen* (Hostettler et al., 2017) to the C-terminal of the *end-1* gene. The CRISPR/Cas9 protocol used to perform knockin was the same as that for knockout except an additional plasmid with a repair template (end-1HR) was constructed and used. The repair template sequence included a ~1000 bp arm homologous to the *end-1* gene sequence on both ends of the *mNeonGreen* coding sequence, and was cloned into the pPD95.77 plasmid between the AgeI and PstI restriction sites. Animals with *mNeonGreen* knockin were detected by PCR and confirmed by sequencing.

### Embryo Mounting

Embryo collection and mounting were performed as previously described (Bao and Murray, 2011). Briefly, 4-6 gravid adult worms with one row of eggs were transferred into a droplet of M9 buffer on a Multitest slide (MP Biomedicals) and cut open under the dissecting microscope (Nikon SMZ745). Two- to four-cell stage embryos were transferred using an aspirator tube assembly (Sigma-Aldrich) into a 2.5 μL droplet of egg buffer containing around thirty 20 μm polystyrene microspheres (Polyscience) on a 24 × 60 mm coverslip (Fisherbrand). Embryo positions were adjusted using an eyelash to produce clusters of two or three embryos, after which an 18 × 18 mm coverslip was laid on top of the droplet, followed by sealing with melted Vaseline. The same procedure was used to mount embryos in an uncompressed manner, except that 45 μm polystyrene microspheres (Polyscience) were used to ensure the spacing between two coverslips was larger than the thickness of embryos (~30 μm).

### 3D Time-lapse Imaging

3D time-lapse imaging was performed using spinning disk confocal microscopy (Revolution XD) with an Olympus IX73 inverted microscope, a Yokogawa CSU-X1 spinning-disk unit, an ASI PZ-2150-XYZ stage with Piezo-Z positioning, and an Andor iXon Ultra 897 EMCCD camera. Images were taken using MetaMorph acquisition software under 60X magnification (PLAPON 60XO N.A. 1.42 objective) with 30 Z slices (1 μm spacing) for six hours at 75-second intervals. The resulting voxel size at each time point was 0.22 × 0.22 × 1.00 μm (X, Y, Z). Imaging parameters (laser power and exposure time) were optimized to minimize photo-damage while maintaining a high signal to noise ratio, and all embryos hatched after long-term imaging.

### Cell Identification, Tracing, and Cell Lineage Construction

4D images were processed with the StarryNite software (Bao et al., 2006; Santella et al., 2014; Santella et al., 2010) for automated cell identification, tracing, and cell lineage reconstruction, and the results were visualized and manually curated using AceTree software (Boyle et al., 2006; Katzman et al., 2018). Detailed procedures for cell identification, lineage tracing, error detection, error correction, and quality control were described in a previous paper (Du et al., 2015). First, a nucleus-localized, ubiquitously expressed mCherry fluorescence signal was used to segment nuclei at each time point using a hybrid blob-detection algorithm (Santella et al., 2010). Second, nuclei were traced over time using a semi-local neighborhood-based framework (Santella et al., 2014). Third, the raw results from automated nuclei identification and tracing were subjected to systematic manual inspection and error correction using AceTree (Boyle et al., 2006; Katzman et al., 2018). AceTree allows visualization of cell lineages and provides a convenient interface for linking lineage tracing results to the original 4D images for efficient detection and correction of errors. Finally, a unique name was assigned to each identified cell according to the Sulston nomenclature (Sulston et al., 1983). Briefly, cell identity (ABa, ABp, EMS, and P2) and body axis were first determined based on the characteristic arrangement of cells in four-cell stage embryo. Then, names were assigned to daughter cells based on the mother cell name and cell position relative to the body axis following cell division. For example, ABal is the daughter cell of ABa that is located on the left side following ABa cell division; ABala is the daughter cell of ABal that is located on the anterior side following ABal cell division. The names of most cells follow this general rule, excepting several early progenitor cells designated with special names (e.g. MS, E, C, D). More information on the naming scheme is given elsewhere (Bao et al., 2006; Santella et al., 2010; Sulston et al., 1983). After lineage tracing, cells present at each time point and associated phenotypic information including cell name; nucleus diameter; X, Y, Z positions; and fluorescence intensity were systematically digitized for further analysis.

### Measurement of Cell Position Noise in Wild-type Embryos

Definition of relative cellular position: We used the 3D coordinates of the centroid of the nucleus as a proxy for cellular position and defined relative cellular position as a vector of geometrical distance from the centroid of the nucleus of the cell of interest (target cell) to that of all other cells (reference cells) present at the same time point (Figure 1C). Because the imaging resolutions are different for XY and Z (pixel size 0.22 vs. 1), the value of Z was multiplied by 4.5 to match the XY scale when calculating geometrical distance. To control for embryo size variation, distances were normalized by the mean distance of all pairwise cell-cell distances in the embryo.

Comparison of relative cellular position: CPN was measured by comparing distance vectors of equivalent cells between wild-type embryos. The root-mean-square deviation (RMSD) values between two distance vectors for all pair-wise comparisons were measured and then averaged to quantify CPN.

Time alignment: Due to variability in cell cycle length, the present time for a specific cell in different embryos could differ. To ensure accurate comparisons of cellular position at a given time, we mapped the absolute time of each cell to a relative scale and calculated CPN at a comparable relative time across all wild-type embryos (Figure S1C). For a given target Cell-X, we first screened the same cell in all 28 embryos to identify the cell with the longest present time and used it as the reference time point for equivalent cells in other embryos. Next, all time points for Cell-X in other embryos were scaled and mapped onto the closest reference time point. The median value of present time for all cells was 32; our definition of comparable time thus on average has a difference less than 1.56% of the total cell cycle length (0.5*1/32). Whenever the CPN of a cell was used for analysis, its CPN at all timepoints was averaged.

Selection of reference cells: Variations in cell cycle length also result in a differential composition of reference cells at a given time point in different embryos. To ensure fair comparison of cell positions, CPN calculations for pair-wise comparisons only included reference cells that were present in both embryos.

### Assessment of the Accuracy and Reliability of Noise Measurement

We validated the accuracy and reliability of CPN measurements in several ways. First, we confirmed that cellular position at each timepoint was accurately determined. Cell identification and tracing were performed automatically using a highly effective algorithm with an accuracy of over 99% (Du et al., 2015). The raw results were further systematically inspected and corrected for residual errors. After the manual curation of cell identification and tracing results in the 28 embryos used for CPN measurements, 125 cells were randomly selected to evaluate the correctness of cellular position determinations. Each cell was manually traced for ten consecutive time points, totaling 1250 instances, to whether cellular position assignments were consistent with the corresponding 3D images. We found all but one instance was correct, leading to an evaluated accuracy of 0.999 (1249/1250).

Second, we validated the robustness of the relative method for defining cellular position as a measure of CPN. We applied this relative approach, quantifying a cell’s position as the distance to reference cells (Figure 1C), in order to circumvent requirements for embryo rotation and positional alignment, which could introduce bias. However, a caveat with this approach is that a cell’s position and noise measurement could be significantly affected by those of its reference cells. We measured CPN levels using subsets (10%, 20%, and 50%) of reference cells and showed that they were highly correlated with those obtained using all reference cells (Figure S1A), confirming that CPN measurement is insensitive to the constituency of reference cells. We also validated that the CPN levels of reference cells have only minor influence on the CPN levels of target cells. We estimated this by computationally altering a cell’s 3D position to introduce *in silico* noise and then determined its impact on CPN measurement. This manipulation caused significantly larger changes in CPN when the cell was treated as a target cell as compared to when it was treated as a reference cell (Figure S1B, Wilcoxon signed-rank test, *p* = 2.28e-112). These results suggest that although the positions of other cells are used as references, the relative method is most sensitive to the CPN of the cell of interest.

Third, we determined the positions of cells across multiple time points and ensured that their CPN levels were measured at comparable times. We aligned cells by the relative time between embryos and compared cellular positions at comparable relative times (Figure S1C). This treatment ensured that temporal measurements of CPN were precise. However, a small-scale time misalignment was inevitable after alignment, due to temporal resolution of the imaging. We modeled the influence of this artifact by artificially misaligning cell times for one time interval (75 seconds, equivalent to 3.4% of the average cell cycle length). We found CPN levels with or without this misalignment were highly correlated (Pearson correlation, R = 0.985, *p* < 2.22e-308), with an average coefficient of variation of 0.01. This excludes the possibility that small-scale time misalignment affects CPN measurement.

Fourth, we confirmed that the number of embryos analyzed was sufficient to represent CPN. We simulated the effects of sample size on CPN levels by sampling smaller numbers of embryos (n = 5, 10, 15, and 20) and comparing to the total 28 embryos. We found that the coefficient of variation of CPN levels was less than 0.05 at an embryo number of ten (Figure S1D). Moreover, we found that CPN levels in embryo sampling replicates were highly correlated (Figure S1E), suggesting CPN is a stable trait of cells.

Finally, we confirmed that the mounting method used for live imaging does not dramatically affect CPN. Two types of mounting methods are widely used for *C. elegans* live imaging, differentiated by whether the embryos are compressed (Bao et al., 2006) or uncompressed (Jelier et al., 2016) during imaging. We found that CPN levels for the two methods were highly correlated (Figure S1F, Spearman rank correlation, ρ = 0.79, *p* = 1.38e-155). However, CPN levels for compressed embryos were significantly lower than that of compressed (Wilcoxon Signed-Rank test, *p* = 7.27e-96), suggesting cellular position is more reproducible when the embryos are compressed. This is consistent with a previous observation that slight compression allows the embryo to rotate to a stereotypical orientation (Sulston et al., 1983).

Collectively, the above analysis of the reproducibility, sensitivity, and robustness of CPN measurement systematically validated the accuracy and reliability of our measurement at high temporal resolution. In this study, we used CPN data from compressed embryos for further analyses.

### Measurement of Cellular Position Noise of Mutant Embryos

*glp-1(e2142)* is a temperature sensitive allele (Priess et al., 1987); these worms were maintained at 15 °C and switched to 25 °C for eight hours before collecting embryos for imaging. *skn-1(zu67)* is a maternal effect recessive allele causing 100% embryonic lethality (Bowerman et al., 1992). We balanced *skn-1(zu67)* using an ubiquitously expressed GFP transgene (oxTi994, Integration site: IV, 2.45) inserted close to the *skn-1* locus (Frokjaer-Jensen et al., 2014) and collected embryos from *skn-1(zu67)* homozygote worms (determined by animals without GFP expression) for imaging. Similarly, because *end-1&ric-7(ok558); end-3(dev85*) mutants have 100% embryonic or larval lethality, that double mutant was balanced by a ubiquitously expressed GFP transgene (oxTi991, Integration site: V, 5.59) inserted near the *end-1* and *end-3* loci (Frokjaer-Jensen et al., 2014). Embryos from heterozygote adult worms were collected for imaging, and embryos that were homozygous for *end-1* and *end-3* (determined by embryos without GFP expression) were used for further analysis.

Measurement of CPN in mutant embryos was performed as in the *wild type* except that for certain mutants, the reference cells used were carefully selected. Some mutations induced widespread, incompletely penetrant changes in cell lineage identity which could cause ambiguity in CPN measurement because the positions of cells with and without the phenotypes are compared. For these cases, we only used reference cells that were deemed to be unaffected. For *glp-1(e2142)* mutants, cells from the ABala sub-lineage were used as reference cells to calculate and compare CPN in both mutants and wild-type embryos (Figure 2D). For *skn-1(zu67)* mutants, cells from the C sub-lineage were used as reference cells (Figure 2E). For *end-1;end-3* double mutants, all cells except gut cells were used as reference cells.

### Identification of Cells with Similar Localization

Hierarchical clustering was applied to the pair-wise cell-cell distance matrix to identify clusters of cells with similar localization using MeV software (Wang et al., 2017) with the following parameters:

Euclidian distance, average linkage clustering, optimize leaf order. A tree cutoff of five was used to identify cell clusters (Figure 3A).

### Quantification of Fate Determinant Expression in Single Cells

Dual color fluorescent reporter strains were constructed to profile single-cell expression of fate determinants during embryogenesis. In each strain, nucleus-localized, ubiquitously expressed mCherry was used to identify and trace nuclei, and GFP fused with the protein sequence of the fate determinant was used to simultaneously quantify protein expression level. Live imaging, cell identification and tracing, cell lineage construction, and quality control were performed as for CPN measurement. All analyzed fate determinants were transcription factors with nuclear localization; this facilitates straightforward quantification of GFP intensity once the nuclei were identified and traced. Quantification of GFP expression within the nucleus was performed using a previously reported method (Murray et al., 2008). Briefly, GFP intensity was first measured within each nucleus at each time point, and then with the local background intensity subtracted. Background intensity was estimated by measuring the GFP intensity within 1.2 r to 2 r from the nuclear centroid for each nucleus (r being the radius of the nucleus). Empirically, an intensity of more than 2,000 A.U. is considered expression. Tree visualizations of singlecell protein expressions were generated by AceTree.

### Identification of Temporal Clusters of CPN Dynamics

Clusters with differential CPN dynamics profiles were identified by applying hierarchical clustering to the time series CPN level for individual cell tracks of leaf cells. To facilitate the comparison of CPN dynamics between cell tracks, CPN levels were normalized to the mean within each cell track. Clustering parameters were: Euclidian distance, average linkage clustering, and a distance cutoff of three to identify clusters (Figure 5C).

### Simulations

Simulation of the sensitivity of CPN measurement: To introduce noise, we randomly changed the 3D position of a cell in one of the 28 embryos by less than 5% of the embryo’s long axis. After 100 simulations, we then calculated the CPN levels of manipulated cells and other unaffected cells to determine the influence of this manipulation (Figure S1B).

Simulation of CPN dynamics: To model CPN dynamics over time (Figure 5B), we simulated the cellular positions for multiple cells (n = 100) in 28 embryos over 100 time points. The initial cellular positions were identical (CPN = 0), and for each time interval we allowed all cells in each embryo to move randomly over a given distance but within the predefined embryonic space. CPN levels at each time point were calculated as for real data. Three distances were modeled, 0.5D, 1D, and 2D, where D is the median distance of cell displacement over one time interval in real embryos.

Simulation of the expected CPN level: To model expected CPN levels, we randomized the direction of cell displacement but kept its distance identical to that of real embryos (Figure 6B and Figure 6C). To simulate the expected CPN level for a cell, we first determined in the real embryo the displacement distance of the cell from its birth time to its division time (Dc). We then allowed the cell at Dc to move in a random direction from the position of its birth time. This simulation was applied to all cells in all embryos. The same strategy was used for simulating the expected CPN level of a time interval, except the cell displacement distance within a time interval (Dt) was used.

### Statistics

All statistical methods and corresponding *p* values were described in the main text or figure legends. Statistical analyses including Pearson correlation, Spearman rank correlation, Mann-Whitney U Test, Mood Median Test, Wilcoxon signed-rank test, *t*-test, and Hypergeometric test were performed using Python and Minitab.

